# Structural basis for late maturation steps of the human mitoribosomal large subunit

**DOI:** 10.1101/2021.03.15.435084

**Authors:** Miriam Cipullo, Genís Valentín Gesé, Anas Khawaja, B. Martin Hällberg, Joanna Rorbach

## Abstract

Mitochondrial ribosomes (mitoribosomes) synthezise a critical set of proteins essential for oxidative phosphorylation. Therefore, their function is vital to cellular energy supply and mitoribosomal defects give rise to a large and diverse group of human diseases ^1^. The architecture of mitoribosomes is strikingly different from that of their bacterial and eukaryotic cytosolic counterparts and display high divergence between species ^2–6^. Mitoribosome biogenesis follows distinct molecular pathways that remain poorly understood. Here, we determined the cryo-EM structures of mitoribosomes isolated from human cell lines with either depleted or overexpressed mitoribosome assembly factor GTPBP5. This allowed us to capture consecutive steps during mitoribosomal large subunit (mt-LSU) biogenesis that involve normally short-lived assembly intermediates. Our structures provide important insights into the last steps of 16S rRNA folding, methylation and peptidyl transferase centre (PTC) completion, which require the coordinated action of nine assembly factors. We show that mammalian-specific MTERF4 contributes to the folding of 16S rRNA, allowing 16S rRNA methylation by MRM2, while GTPBP5 and NSUN4 promote fine-tuning rRNA rearrangments leading to PTC formation. Moreover, our data reveal an unexpected role for the elongation factor mtEF-Tu in mt-LSU assembly, in which mt-EF-Tu interacts with GTPBP5 in a manner similar to its interaction with tRNA during translational elongation. Together, our approaches provide detailed understanding of the last stages of mt-LSU biogenesis that are unique to mammalian mitochondria.

## Main

Mammalian mitoribosomes assemble in a multi-step process that includes the maturation of two ribosomal RNAs (rRNAs; 12S and 16S), a structural tRNA, and incorporation of 82 mitoribosomal proteins (MRPs) ^7^. Multiple assembly factors, many specific to mammalian mitochondria, assist in the mitoribosome assembly process. A growing number of studies in recent years have shown that a family of GTP-binding proteins (GTPBPs) is crucial for mammalian mitoribosome assembly ^8–13^. Among these GTPBPs, GTPBP5 participates in the late steps of large subunit (mt-LSU) maturation, and its deletion leads to severe translational defects ^8,11^.

To understand the molecular basis for the late stages of human mitochondrial mt-LSU assembly, we used single-particle electron cryomicroscopy (cryo-EM) to determine the structure of an mt-LSU intermediate isolated from GTPBP5-deficient cells (GTPBP5^KO^). In addition, we determined the cryo-EM structure of a GTPBP5-bound mt-LSU intermediate immunoprecipitated from cells expressing a tagged variant of GTPBP5 (GTPBP5^IP^).

### Composition of the GTPBP5^KO^ and GTPBP5^IP^ mt-LSU assembly intermediates

Both the GTPBP5^KO^ and GTPBP5^IP^ mt-LSU assembly-intermediates reveal several trapped assembly factors: the MTERF4-NSUN4 complex, MRM2, MTG1 and the MALSU1:L0R8F8:mt-ACP module (Fig. 1a,b, Extended Data Fig. 1 and 2). Furthermore, the GTPBP5^IP^ mt-LSU structure features GTPBP5 and the mitochondrial elongation factor mtEF-Tu (Fig. 1b, Extended Data Fig. 2). Comparing the GTPBP5^KO^ and the GTPBP5^IP^ mt-LSU intermediates with the mature mt-LSU ^14^ reveals two crucial differences in the 16S rRNA conformation (Fig. 1c,d). First, in both GTPBP5^KO^ and GTPBP5^IP^ intermediates, MTERF4 in the MTERF4-NSUN4 complex, binds an immaturely folded region of the 16S domain IV. This region (C2548-G2631) – corresponding to helices H68, H69, and H71 of the mature mt-LSU – is folded into a novel intermediate rRNA helical structure, hereafter denoted helix-X (Fig. 1). The helix-X occupies a different position on the mt-LSU than H68-71 in the mature mt-LSU, where helices H68-71 and H89-90 jointly form the peptidyl-transferase centre (PTC) (Fig. 1d). MTERF4s binding of helix-X partly orders the disordered rRNA in the mt-LSU assemblies’ subunit interface side (Extended Data Fig. 1) and thereby enables MRM2 to bind (Fig. 1d). Second, in the GTPBP5^KO^ – but not in the GTPBP5^IP^ – the junction between H89 and H90 of domain V is significantly different compared to the mature mitoribosome (Fig. 1c, d). Specifically, at the base of H89, one helical turn remains unfolded and instead forms a flexible loop in the GTPBP5^KO^ (Fig. 1d).

**Fig. 1:**
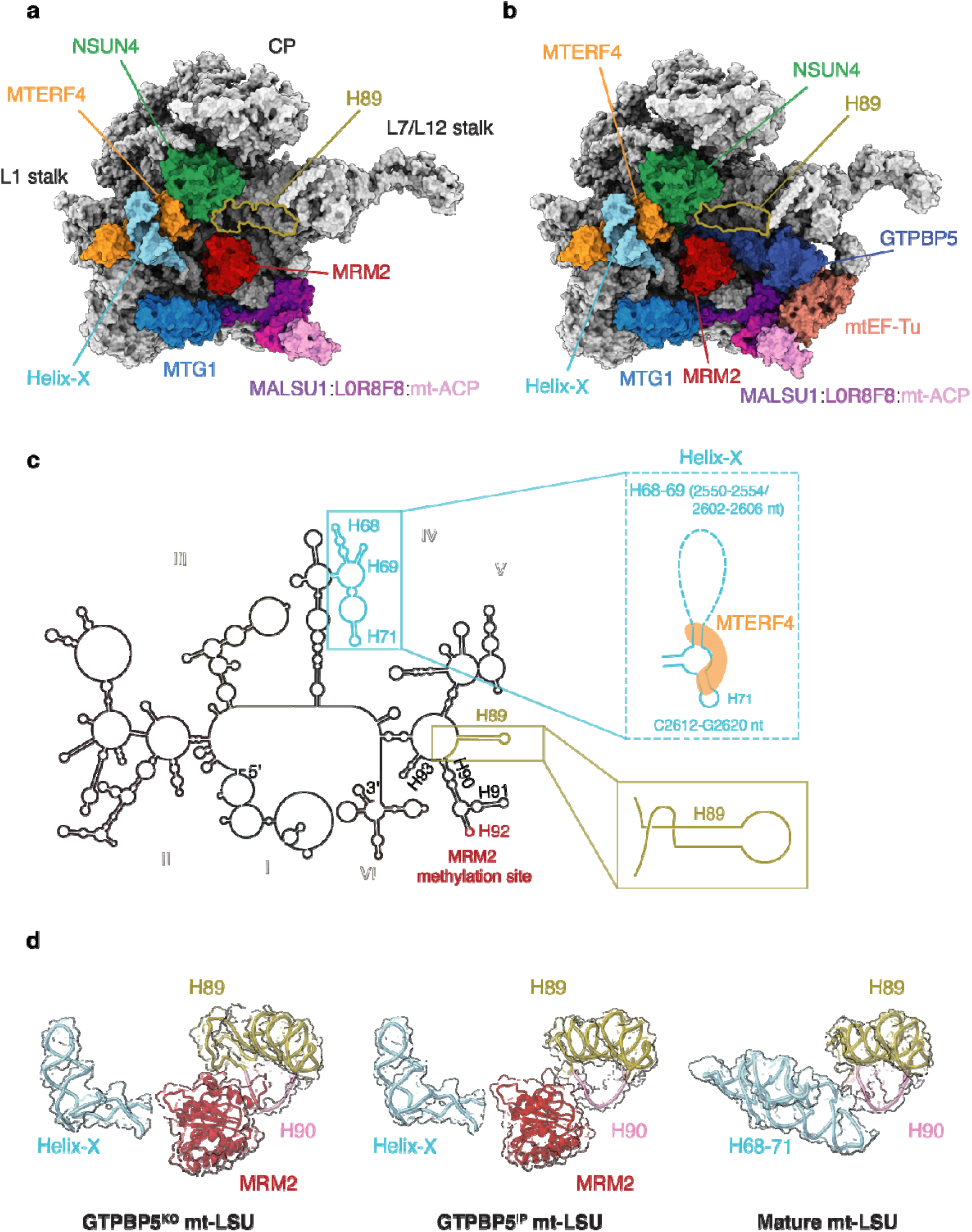
Overview of the GTPBP5^KO^ and the GTPBP5^IP^ mt-LSU assembly intermediates and comparison with the mature mt-LSU. **a**, The GTPBP5^KO^ is bound by MTERF4, NSUN4, MRM2, MTG1 and the MALSU1 module. Mitoribosomal proteins and 16S mt-rRNA are shown in grey. Helix-X bound to MTERF4 is highlighted as well as H89. **b**, The interface of the GTPBP5^IP^ mt-LSU intermediate associated with MTERF4, NSUN4, MRM2, MTG1, MALSU1:L0R8F8:mt-ACP complex, GTPBP5 and mtEF-Tu. Helix-X bound to MTERF4 is shown in light blue. **c**, Secondary structure of the mature mt-LSU 16S mt-rRNA. Differences in the rRNA fold of the GTPBP5^KO^ mt-LSU intermediate are shown in the zoomed-in views. Dashed lines indicate regions that are not modelled. MRM2 methylation site (H92) is indicated in red. The six 16S mt-rRNA domains are shown in different colours. **d**, Positioning of helix-X (H68-71) and helices H89 and H90 in GTPBP5^KO^ mt-LSU (left), GTPBP5^IP^ mt-LSU (middle) and the mature mt-LSU (right) (PDB:6ZSG ^15^). In the GTPBP5^KO^ and the GTPBP5^IP^ mt-LSU structures MRM2 is present.

### MTERF4-NSUN4 complex steers the final steps of 16S rRNA folding and allows for MRM2 binding

The MTERF4-NSUN4 complex, previously shown to be essential for monosome assembly ^16,17^, binds at the intersubunit interface in our GTPBP5^KO^ and GTPBP5^IP^ structures (Fig. 1a, b). The C-terminal part of MTERF4 binds to NSUN4 close to the NSUN4 N-terminus in a mixed hydrophobic-polar binding interface similar to earlier crystal structures of the isolated complex ^18,19^ (Fig. 2a). NSUN4 was previously shown to m^5^C-methylate the C1488 carbon 5 in 12S mt-rRNA ^17^. In our structures, the active site of NSUN4 is turned towards the mt-LSU core (Extended Data Fig. 3a), impeding methylation of the 12S mt-rRNA. Although the methyl-donor S-adenosyl-methionine (SAM) is observed in the NSUN4 active site, no RNA substrate is present. Furthermore, in the GTPBP5^KO^ and GTPBP5^IP^ structures, the MTERF4-NSUN4 complex is bound and bent from two sides by uL2m. Specifically, a uL2m C-terminal extension penetrates in between NSUN4 and MTERF4 to further stabilize the MTERF4-NSUN4 binding interface and decreases the curvature of the MTERF4 solenoid relative to the crystal structures (Fig. 2a). This reforming of the MTERF4 solenoid is necessary to bind the helix-X rRNA region in the strongly positively charged concave side of MTERF4 (Extended Data Fig. 3b). Here, MTERF4 forms an extensive network of contacts with helix-X that stabilizes the association and promotes helix-X folding (Fig. 2b). The mature H71 base-pairing is already formed within helix-X. Thus, by binding to helix-X, MTERF4 intiates the folding of this 16S mt-rRNA region. Furthermore, it also exposes the A-loop, which is obstructed by H68, H69, and H71 in the mature mt-LSU (Fig. 1d), thereby allowing MRM2 binding.

**Fig. 2:**
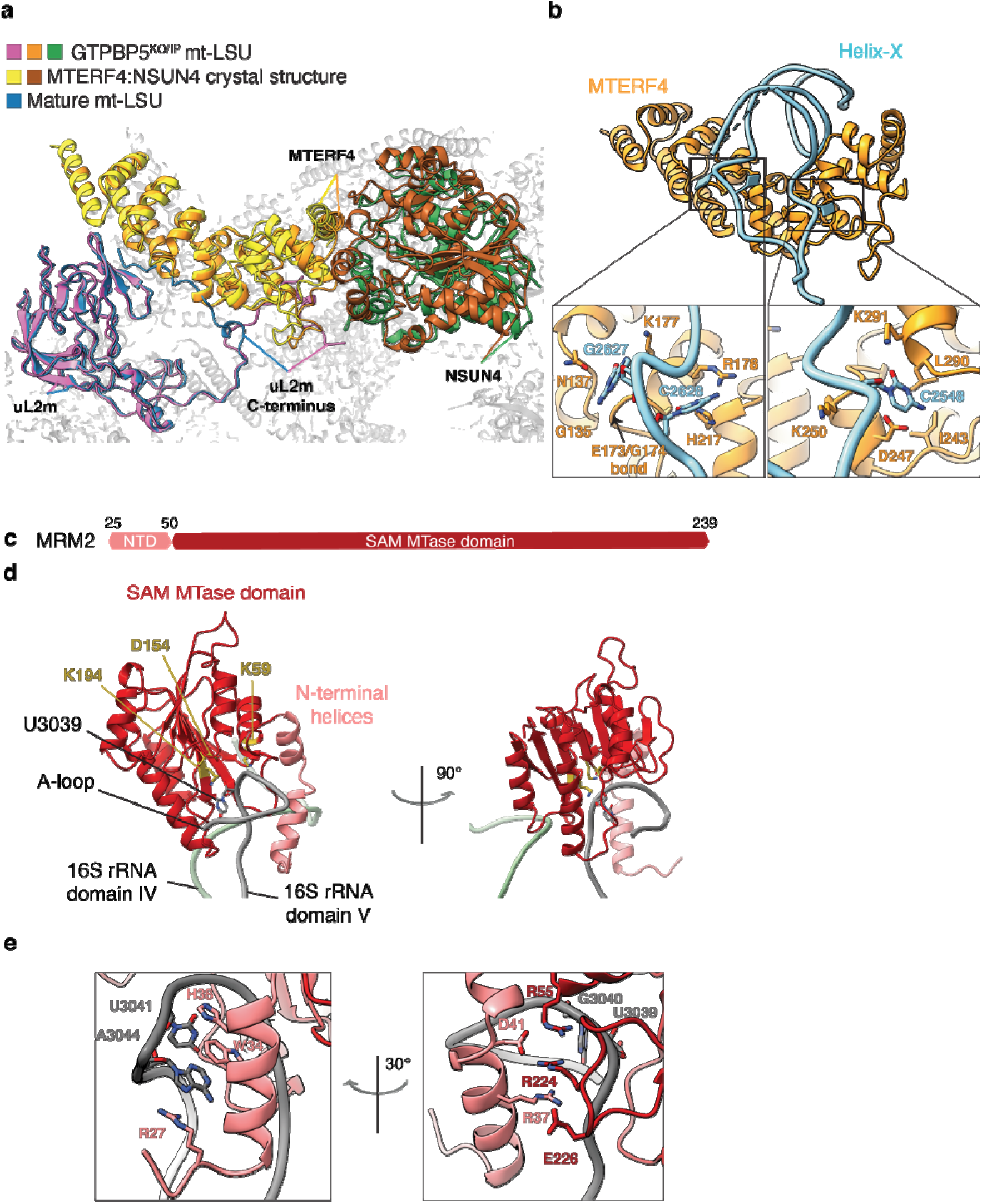
MTERF4-NSUN4 and MRM2 interaction with the mt-LSU assembly intermediates. **a**, Comparison of the MTERF4-NSUN4 complex bound to the GTPBP5^KO/IP^ mt-LSU (orange and green respectively) with the MTERF4-NSUN4 crystal structure (PDB: 4FP9 ^18^) (yellow and brown, respectively), and of uL2m from the GTPBP5^KO/IP^ mt-LSU (pink) with uL2m from the mature mt-LSU (blue) (PDB: 3J7Y ^14^). The uL2m C-terminus is indicated in both structures. Helix-X is not shown. **b**, MTERF4-NSUN4 complex bound to helix-X. Zoom-in panels show the interactions of MTERF4 with helix-X. **c**, Schematic representation of MRM2 domains (NTD - light pink, SAM MTase domain - red). **d**, MRM2 interaction with the domain IV rRNA (nucleotides 2644-2652, green) and the A-loop (grey). The MRM2 methylation site (U3039) as well as the catalytic triad of MRM2 (K59, D154, K194) are highlighted as sticks. **e**, Zoomed-in views showing MRM2 interactions with the A-loop in different orientations.

Similarly to a previously determined mt-LSU assembly intermediate ^20^ there is a MALSU1-module positioned adjacent to uL14m in both the GTPBP5^KO^ and GTPBP5^IP^ (Fig.1a,b). Furthermore, MTG1 (GTPBP7), which assists in late-stage mt-LSU maturation ^21^, is bound in the vicinity of helix-X (Fig. 1a,b). MTG1 contacts the C-terminus of MALSU1 (Extended Data Fig. 4a) and the region encompassing A2554-U2602 of helix-X. This region could not modelled due to the lower local resolution, but the contact is visible in the electron density map (Extended Data Fig. 4b). Interestingly, the position of human MTG1 in our structures differs significantly from its bacterial and trypanosomal counterparts (^3,22^, Extended Data Fig. 4c). Specifically, while in other systems MTG1 homologs contact the rRNA, reaching out towards the PTC (Extended Data Fig. 4c), in the trapped intermediates described here MTG1 is unlikely to induce pronounced conformational changes of the PTC or participate in the recruitment/dissociation of assembly factors.

MRM2 2′-O-methylates U3039 in the 16S A-loop during mt-LSU assembly ^23,24^ and in our GTPBP5^KO^ and GTPBP5^IP^ structures, MRM2 binds in the mt-LSU intersubunit interface (Fig. 1a,b). It features two N-terminal α-helices followed by a canonical S-adenosyl-L-methionine-dependent methyltransferase domain (SAM MTase) (Fig. 2c). In GTPBP5^KO^, but not in GTPBP5^IP^, the two N-terminal α-helices extend from MRM2 and insert into the rRNA core to thereby displace and retrieve the A-loop (16S mt-rRNA domain V) through a complex interaction network (Fig. 2d,e). This places the 2′-hydroxyl of U3039 close to the ideal methyl-acceptor position in the MRM2 active site (Fig. 2d). However, there is no density for either SAM or S-adenosyl homocysteine (SAH) in the MRM2 active site and there is no apparent density for a 2′-*O*-methyl on U3039 (Extended Data Fig. 5). Interestingly, G3040 that is 2′-*O*-methylated by MRM3 ^23,24^, is methylated in our structures (Extended Data Fig. 5). Hence, 2′-O-methylation by MRM3 takes place prior to MRM2 methylation in human mitoribosome biogenesis.

### GTPBP5 promotes remodelling of the PTC

GTPBP5 consists of a glycine-rich N-terminal domain (Obg-domain) and a C-terminal GTPase domain (G-domain) (Fig. 3a). In our GTPBP5^IP^ structure, the G-domain has GTP in its active site and is wedged between the L7/L12 stalk and the MALSU1 module (Fig. 1b and Extended Data Fig. 6a). The Obg-domain protrudes into the PTC (Fig. 3b), thereby displacing the A-loop from the MRM2 active site and expelling the MRM2 N-terminal α-helices from the rRNA core (Fig. 3c), while the A-loop folds into the fully mature position (Fig. 3b3).

**Fig. 3:**
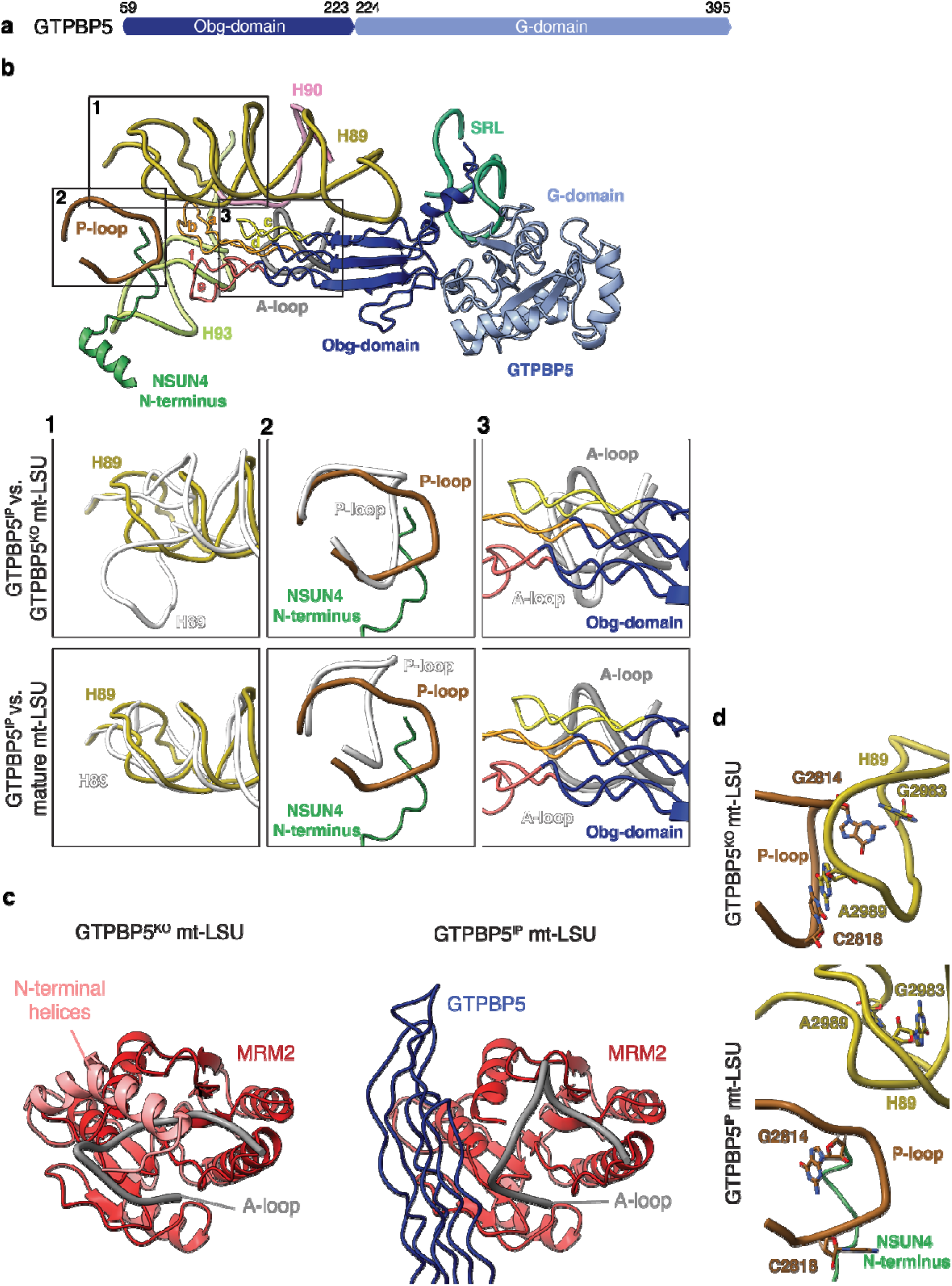
GTPBP5 contributes to the maturation of the PTC region. **a**, Schematic representation of GTPBP5 domains (Obg-domain dark blue, G-domain light blue). **b**, Overview of GTPBP5 interactions with the 16S rRNA. The Obg-domain (dark blue) contacts helices that are in the PTC region: P-loop, A-loop, H89, H90, H93. Helices a-f of GTPBP5 Obg-domain are indicated. The SRL and the NSUN4 N-terminus are shown. Boxes 1-3 show the remodelling of the PTC in GTPBP5^IP^ mt-LSU (in color) compared with GTPBP5^KO^ mt-LSU (in white, higher panel) and with the mature mt-LSU (in white, lower panel) (PDB: 6ZSG^15^). **c**, Comparison of MRM2 (red) and the A-loop (grey) conformations between GTPBP5^KO^ mt-LSU (left) and GTPBP5^IP^ mt-LSU (right). The N-terminal helices (pink) of MRM2 could not be modelled in the GTPBP5^IP^ mt-LSU. The GTPBP5 Obg-domain is shown in dark blue. **d**, Comparison of the P-loop and H89 conformations between GTPBP5^IP^ mt-LSU (lower panel) and GTPBP5^KO^ mt-LSU structures (higher panel).

The protruding Obg-domain is positioned between H89 and H93 and occupies the space that accommodates the acceptor arm of the A-site tRNA during translation (Extended Data Fig. 6b). Hereby, GTPBP5 adopts a tRNA mimicry strategy, similar to ObgE of *E. coli* ^25^. The Obg-domain contains six glycine-rich sequence motifs that form antiparallel polyproline-II helices (helices a–f) (Fig. 3b). Helices c and d bind the A-loop, while the loop between helices e and f inserts into the major groove of H93. The loop between a and b inserts at the triple-junction formed between H89-H90-H93 (Fig. 3b).

Comparison of the GTPBP5^KO^ and the GTPBP5^IP^ structures with the mature mt-LSU reveals extensive maturation of the PTC upon GTPBP5 binding. The partly unfolded H89 in the GTPBP5^KO^ is folded in the GTPBP5^IP^ (Fig. 3 b1). This folding is coordinated by the joint action of GTPBP5 and NSUN4: in the presence of GTPBP5, the extended N-terminal region of NSUN4 inserts into the rRNA core and temporarily displaces the P-loop (Fig. 3 b2), thereby breaking the P-loop interaction with H89 (Fig. 3d). As a consequence, H89 is given the space necessary to fold into a structure similar to its mature form (Fig. 3 b1, lower).

The GTPBP5^IP^ structure shows a rotation of the L7/L12 stalk in comparison to the GTPBP5^KO^ structure (Extended Data Fig. 6c). Here, the rRNA in the L54/L11 region of the stalk forms π-stacking interactions with two residues of the GTP-ase switch I element of GTPBP5 (Extended Data Fig. 6a,c). Thereby, the L7/L12 stalk stabilizes the “state 2” conformation of the switch I. In this way, the rotated L7/L12 stalk stabilizes the GTP-state of GTPBP5^26^ and consequently a GTP is bound in our structure (Extended Data Fig. 6a,c). The requirement for GTPBP5 to be in a GTP-bound state is supported by the inability of a GTPBP5 Walker A mutant (GTPBP5-S238A) to bind mt-LSU intermediates ^8^. A back-rotation of the L7/12 stalk, presumably by binding of another maturation factor to the mt-LSU assembly intermediate, would lead to a release of the switch I and the activation of GTP hydrolysis, followed by release of GTPBP5 from the mt-LSU assembly intermediate. Taken together, GTPBP5 plays a direct and active role in rRNA remodeling and, together with the NSUN4 N-terminus, orchestrates the maturation of mitoribosomal PTC.

### Translation elongation factor mtEF-Tu is involved in mitoribosome assembly

mtEF-Tu consists of a GTPase domain (G-domain; domain I) and two structurally similar β-stranded domains (domains II and III) (Fig. 4a). It was recently shown that during translation, mtEF-Tu·GTP delivers aminoacylated-tRNA to the mitoribosome in a manner similar to its bacterial EF-Tu counterparts (Extended Data Fig. 7a,^27^). In contrast, the binding of an EF-Tu·GTP·aa-tRNA complex is sterically hindered by the MALSU1 module bound in our mt-LSU intermediates (Extended Data Fig. 7a). Unexpectedly, mtEF-Tu binds to the mitoribosome in a unique manner in our GTPBP5^IP^ structure (Fig. 4b). Here, domains II and III establish extensive interactions with GTPBP5, the sarcin-ricin loop (SRL) and the MALSU1 stalk (Fig. 4b,c). In addition, the G-domain switch I element, in its “state 1”/GDP conformation (Extended Data Fig. 7b), extends and binds MALSU1. Thereby, mtEF-Tu, together with the SRL and MALSU1, forms a platform for GTPBP5 binding (Fig. 4b). These structural conclusions are supported by earlier mass-spectrometry data on isolated GTPBP5^IP^ assembly intermediates and protein-proximity interactome analysis ^8,28^.

**Fig. 4:**
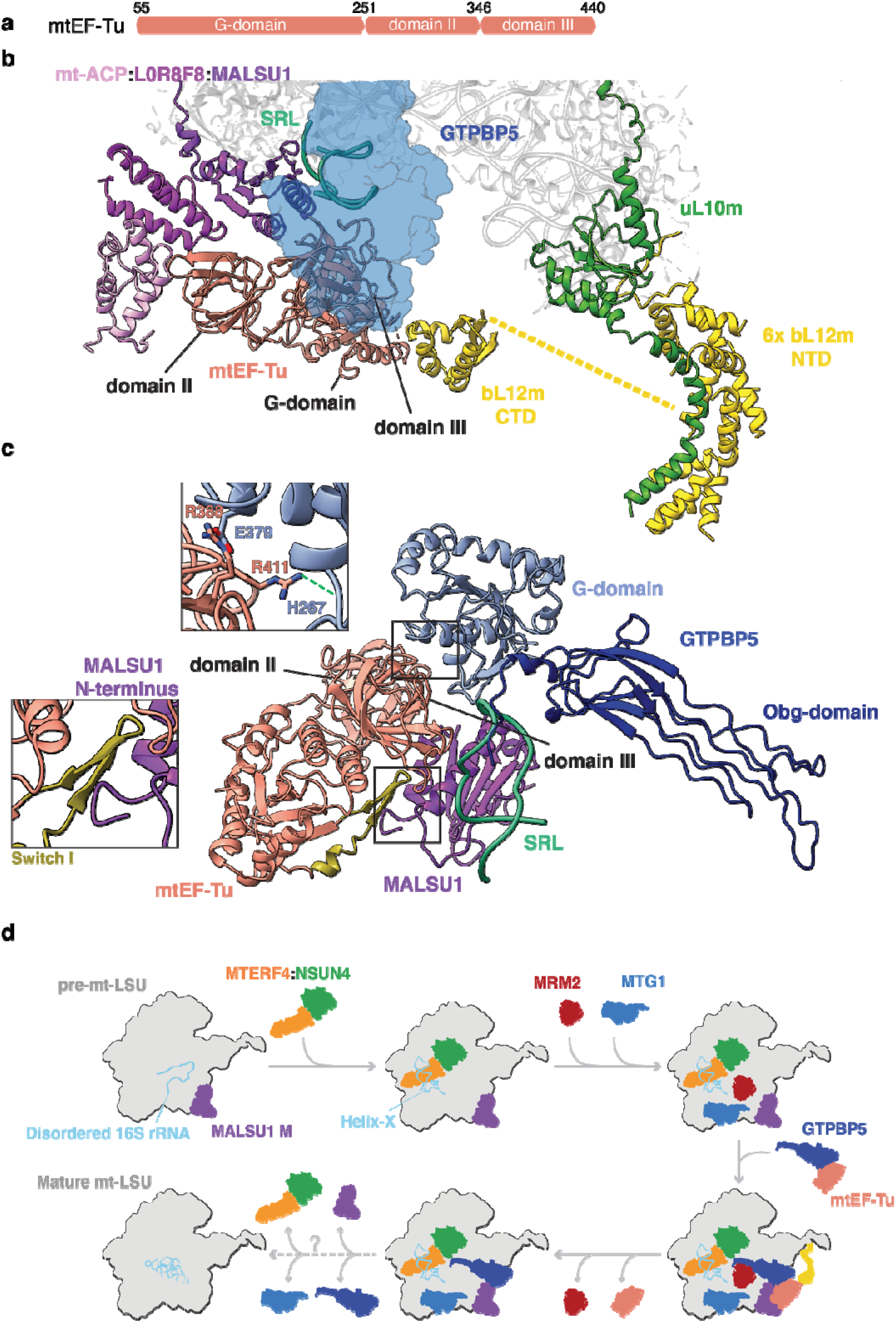
Interaction of mtEF-Tu with the mt-LSU assembly intermediate and model of the final steps of mt-LSU biogenesis. **a**, Schematic representation of mtEF-Tu domains. **b**, mtEF-Tu interaction with GTPBP5, the MALSU1 module and the bL12m C-terminal domain. mtEF-Tu G-domain, domain II and domain III and the SRL are indicated. The six copies of bL12m N-terminal domain and uL10m are also highlighted. The yellow dashed line indicates a hypothetical connection between bL12m CTD and one of the six copies of bL12m NTD, not visible in the structure. **c**, Representation of the mtEF-Tu interaction with GTPBP5 and MALSU1. The upper zoomed-in panel features interactions between the GTPBP5 G-domain and the mtEF-Tu domain III. The green dashed line indicates interactions to the RNA phosphate backbone. The lower zoomed-in panel shows the mtEF-Tu switch I interaction with MALSU1. **d**, Final steps of the mt-LSU assembly. The dashed arrow indicates that biogenesis factors are released in an unknown order.

The G-domain of mtEF-Tu does not contact the SRL as in mtEF-Tu’s canonical role in translation but instead binds to the C-terminal region of a bL12m that also contacts uL10m at the stalk base (Fig. 4b). In bacteria, homologs to bL12m and uL10m, recruit and activate translational GTPases such as EF-Tu via the bL12m C-terminal domain^29,30^ and stimulate GTP hydrolysis 1000-fold^31^. Taken together, this suggests that mtEF-Tu hydrolysis – stimulated by bL12m and uL10m – is used to accommodate GTPBP5 on the maturating mt-LSU in analogy to the canonical EF-Tu role in translation, in which aminoacylated-tRNA is accommodated on the translating ribosome (Extended Data Fig. 7a).

### Concluding remarks

Our analyses shed new light into mammalian mitoribosome maturation and explain the essential roles □ of several assembly factors that together promote fine RNA rearrangments and lead to the mt-LSU completion. Thanks to our approaches that combine biochemical tools with structural determination, we were able to uncovered several features unique to mammalian mitochondria. Based on these data, we propose a model of the late-stage mt-LSU assembly that requires the interplay of nine auxiliary factors (Fig. 4d).

Lastly, as defects in mitoribosome biogenesis – resulting from, for example, mutations in MRM2 and GTPBP5 – are increasingly implicated in mitochondrial disease ^32,33^, the current work does not only describes fundamental cellular processes but may also further new diagnostic and therapeutic approaches to mitochondrial diseases.

## Methods

### Generation of GTPBP5 knock-out cell line

The knock-out cell line (GTPBP5^KO^) was generated in the Flp-In T-Rex human embryonic kidney 293 (HEK293T) cell line (Invitrogen) using CRISPR/Cas9 technology targeted on exon 1 of *MTG2* gene, which encodes for GTPBP5, as described ^8^. In short, two pairs of gRNAs were designed and cloned into the pSpCas9(BB)-2A-Puro (pX459) V2.0 vector to generate out-of-frame deletions. Transfection of HEK293T cell line with the pX459 variants was performed using Lipofectamine 3000 following manufacturer’s instructions. Selection of transfected cells was done using puromycin treatment at a final concentration of 1.5 mg/ml for 48 hours. Subsequently, cells were single-cell diluted and transferred into a 96-well plate. Selected clones were screened via Sanger sequencing and Western blotting.

### Purification of the mt-LSU from GTPBP5^KO^ cell line via sucrose gradient centrifugation

Isolation of mitochondria was performed from GTPBP5^KO^ cell line as described in Rorbach *et al*. ^24^, with some modifications. Crude mitochondria were further purified via differential centrifugation by being loaded onto a sucrose gradient (1 M and 1.5□M sucrose, 20□mM Tris-HCl pH 7.5, 1□mM EDTA) and centrifuged at 25000 rpm for 1 hour at 4°C (Beckman Coulter SW41-Ti rotor). Mitochondria forming a band at the interphase between the 1 M and 1.5 M sucrose were collected and resuspended in 10 mM Tris-HCl pH = 7.5 in 1:1 ratio. After centrifugation, the final mitochondrial pellet was resuspended in mitochondrial freezing bufRNase inhibitor (Invitrogen)) andfer (200□mM trehalose, 10□mM Tris-HCl pH 7.5, 10□mM KCl, 0.1% BSA, 1□mM EDTA), snap frozen in liquid nitrogen and stored at −80°C.

The mt-LSU was purified from the GTPBP5^KO^ cell line via a sucrose gradient centrifugation experiment. Mitochondria were lysed at 4°C for 20 minutes (25 mM HEPES-KOH pH = 7.5, 20 mM Mg(OAc)_2_, 100 mM KCl, 2% (v/v) Triton X-100, 2 mM dithiothreitol (DTT), 1x cOmplete EDTA-free protease inhibitor cocktail (Roche), 40 U/µl RNase inhibitor (Invitrogen)) and later centrifuged at 13000 rpm for 5 minutes at 4°C. For mitoribosome purification, the mitolysate was subjected to sucrose cushion ultracentrifugation method (0.6 M sucrose, 25 mM HEPES-KOH pH = 7.5, 10 mM Mg(OAc)_2_, 50 mM KCl, 0.5% (v/v) Triton X-100, 2 mM DTT) by being centrifuged at 73000 rpm for 45 minutes at 4°C (Beckman Coulter TL120.2 rotor). The mitoribosomal pellet was subsequently resuspended in ribosome resuspension buffer (25 mM HEPES-KOH pH = 7.5, 10 mM Mg(OAc)_2_, 50 mM KCl, 0.05% DDM, 2 mM DTT) and centrifuged at 13000 rpm for 10 minutes at 4°C. The obtained supernatant was then loaded onto a linear sucrose gradient (15-30% (w/v)) in 1x gradient buffer (25 mM HEPES-KOH pH = 7.5, 10 mM Mg(OAc)_2_, 50 mM KCl, 0.05% DDM, 2 mM DTT) and centrifuged for 2 hours and 15 minutes at 39000 rpm at 4°C (Beckman Coulter TLS55 rotor). Fractions corresponding to the large mitochondrial subunit were collected and subjected to buffer exchange (25 mM HEPES-KOH pH = 7.5, 10 mM Mg(OAc)_2_, 50 mM KCl) using Vivaspin 500 centrifugal concentrators.

### Generation of a mammalian cell line expressing GTPBP5

A stable mammalian cell line overexpressing C-terminal FLAG-tagged GTPBP5 (GTPBP5::FLAG) in a doxycycline-inducible dose-dependent manner was generated as described in Cipullo et al. ^8^. The GTPBP5 cDNA (hORFeome Database; Internal ID: 12579) was cloned into pcDNA5/FRT/TO. Flp-In T-Rex human embryonic kidney 293 (HEK293T, Invitrogen) cells were cultured in DMEM (Dulbecco’s modified eagle medium) containing 10% (v/v) tetracycline-free fetal bovine serum (FBS), 2 mM Glutamax (Gibco), 1x Penicillin/Streptomycin (Gibco), 50 µmg/ml uridine, 10 µg/ml Zeocin (Invitrogen) and 100 µg/ml blasticidin (Gibco) at 37 °C under 5% CO_2_ atmosphere. Cells were seeded in a 6-well plate, grown in medium without antibiotics and co-transfected with pcDNA5/FRT/TO-GTPBP5::FLAG and pOG44 using Lipofectamine 3000 according to manufacturer’s recommendations. After 48 hours, selection of cells was promoted by addition of hygromycin (100 µg/ml, Invitrogen) and blasticidin (100 µg/ml) to culture media. After two to three weeks post-transfection, single colonies were picked and GTPBP5 overexpression was tested via Western Blot analysis 48 hours after induction with 50 ng/ml doxycycline.

### Immunoprecipitation experiment

Isolation and purification of mitochondria from GTPBP5::FLAG overexpressing cell line was performed as described in the above paragraph “Purification of the mt-LSU from GTPBP5^KO^ cell line via sucrose gradient centrifugation”. The mt-LSU bound with GTPBP5 was isolated via FLAG-immunoprecipitation analysis (IP). Pelleted mitochondria were lysed at 4°C for 20 minutes (25 mM HEPES-KOH pH = 7.5, 20 mM Mg(OAc)_2_, 100 mM KCl, 2% (v/v) Triton X-100, 0.2 mM DTT, 1x cOmplete EDTA-free protease inhibitor cocktail (Roche), 40 U/□l RNase inhibitor (Invitrogen)) and centrifuged at 5000g for 5 minutes at 4°C. The supernatant was then added to ANTI-FLAG M2-Agarose Affinity Gel (Sigma-Aldrich) previously equilibrated (25 mM HEPES-KOH pH = 7.5, 5 mM Mg(OAc)_2_, 100 mM KCl, 0.05% DDM) and incubated for 3 hours at 4°C. After incubation, the sample was centrifuged at 5000g for 1 minute at 4°C, the supernatant was removed and the gel was washed three times with wash buffer. Elution (25 mM HEPES-KOH pH = 7.5, 5 mM Mg(OAc)_2_, 100 mM KCl, 0.05% DDM, 2 mM DTT) was performed using 3x FLAG Peptide (Sigma-Aldrich) for about 40 minutes at 4°C.

### Cryo-EM data acquisition and image processing

Prior to cryo-EM grid preparation, grids were glow-discharged with 20 mA for 30 seconds using a PELCO easiGlow glow-discharge unit. Quantifoil Cu 300 mesh (R 2/2 geometry; Quantifoil Micro Tools GMBH) covered with a thin layer of 3 nm carbon were used for the for the GTPBP5^KO^ sample. Carbon lacey films (400 mesh Cu grid; Agar Scientific) mounted with ultrathin carbon support film were used for the GTPBP5^IP^ sample. Three □l aliquots of sample were applied to the grids, which were then vitrified in a Vitrobot Mk IV (Thermo Fisher Scientific) at 4°C and 100% humidity (blot 10 s, blot force 3, 595 filter paper (Ted Pella Inc.)). Cryo-EM data collection (Extended Data Table 1) was performed with EPU (Thermo Fisher Scientific) using a Krios G3i transmission-electron microscope (Thermo Fisher Scientific) operated at 300 kV in the Karolinska Institutet’s 3D-EM facility. Images were acquired in nanoprobe EFTEM SA mode with a slit width of 10 eV using a K3 Bioquantum during 1 second during which 60 movie frames were collected with a flux of 0.82 e^−^/Å^2^ per frame. Motion correction, CTF-estimation, Fourier binning (to 1.02 Å/px), picking and extraction in 600 pixel boxes (size threshold 300 Å, distance threshold 20 Å, using the pretrained BoxNet2Mask_20180918 model) were performed on the fly using Warp ^34^. Only particles from micrographs with an estimated resolution of 3.6 Å and underfocus between 0.2 and 3 □m were retained for further processing.

For the GTPBP5^KO^ dataset, 704720 particles were picked from 37307 micrographs (Extended Data Fig. 1). The particles were imported into CryoSPARC 2.15 ^35^ for further processing. After 2D classification, 130289 particles were selected for an ab-initio reconstruction. This reconstruction, in addition to two “bad” reconstructions created from bad 2D class-averages, were used for heterogeneous refinement of the complete particle set resulting in one of the three classes yielding a large-subunit reconstruction with high resolution features (196318 particles). After homogeneous refinement of these particles, the PDB model of a mitochondrial LSU assembly intermediate (PDB: 5OOL ^20^) was fitted in the density. The reconstruction contained the MALSU1 module and also featured weak unexplained densities for several additional components in the intersubunit interface. A 3D variability analysis was performed with a mask on the intersubunit interface and a low pass resolution of 10 Å, and subsequently used for clustering into six particle classes representing different assembly intermediates. Two of the classes (43057 and 41619 particles) lacked the density for the A- and P-loops, H89, helices 68-71 and the L7/12 stalk. The A- and P-loops become visible in the third class (28001 particles). The fourth class revealed a number of biogenesis factors: MRM2, MTERF4-NSUN4, MTG1 and the structured H67-H71 rRNA region (helix-X) (48646 particles) as well as H89. All the biogenesis factors are absent in the fifth class, in which helices 68 and 71 move to the mature position (26678 particles). H69 is nevertheless not visible. The last class contains the small subunit (8317 particles). Non-uniform refinement of the fourth particle set yielded a reconstruction at 2.64. Å, which was used for model building and refinement. As the density for MTG1 was weaker than for the other factors, 3D variability analysis was performed with a mask on the MTG1 region and a 10 Å low-pass filter to select particles containing MTG1 (19254 particles, which was subsequently subjected to homogeneous refinement yielding a reconstruction at 2.90 Å).

For the GTPBP5^IP^ dataset, 283598 particles were picked from 112076 micrographs using WARP and imported into CryoSPARC 2.15 (Extended Data Fig. 2) ^35^. The complete particle set was used in heterogeneous refinement against the same three references derived from the GTPBP5^KO^ dataset. One of the classes (78306 particles) yielded a high-resolution reconstruction of the mt-LSU assembly intermediate. After homogeneous refinement, additional density for GTPBP5 was visible in the intersubunit interface. 3D variability analysis was performed with a mask on the GTPBP5 region and a low pass resolution of 10 Å. Subsequent clustering into two particle clusters revealed a particle subset containing GTPBP5 (71834 particles), which was used for model building and refinement. This reconstruction also features densities for MRM2 and MTERF4-NSUN4. In addition, a weak density was present for mtEF-TU, the bL12m C-terminal domain and MTG1. The refined particles were subject to 2D classification and the bad classes were removed. The remaining particles were polished and refined in Relion 3.1 and re-imported into CryoSPARC for further processing. 3D variability analysis was performed on these particles with a mask covering mtEF-TU, bL12m and MTG1 and a 10 Å low pass filter. Subsequent clustering (4 clusters) revealed 2 clusters containing mtEF-TU/bL12m (17886 particles in total), one cluster containing MTG1 (8233 particles) and one cluster containing all three proteins (13376 particles). The reconstructions derived from the MTG1- and the mtEF-Tu-containing particles reached a resolution of 3.19 and 3.21 Å respectively.

### Model building and refinement

Model building of the GTPBP5^KO^ and GTPBP5^IP^ mt-LSU assembly intermediate structures was performed using *Coot* ^36^. The structure of a previous mt-LSU assembly intermediate (PDB 5OOL, ^20^) was used as a starting model. MRM2 and MTERF4-NSUN4 were identified by modelling secondary structure elements in *Coot*, and using the initial models for a structural search using the DALI server ^37^. MTG1, as well as mtEF-Tu and the bL12m C-terminal domain in the GTPBP5-bound mt-LSU dataset, were identified using a density-based fold-recognition pipeline ^20^. Using SWISS-MODEL ^38^, we generated homology models for the human GTPBP5 (template: PDB 4CSU chain 9 ^25^), MTG1 (template: PDB 3CNL chain A ^39^), bL12m (template: PDB 1DD3 chain A ^40^) and mtEF-TU (template: PDB 1D2E chain A ^41^). All the models, as well as the crystal structure of the human MTERF4-NSUN4 (template: PDB 4FP9 chains A and B ^18^), were fitted into the density map using *Coot* JiggleFit. The MTG1 GTPase domain and the L17/12 stalk were excluded from atomic refinement and were only subject to rigid body refinement. Metal ions and modifications were placed based on map densities. Stereochemical refinement was performed using PHENIX ^42^. Refinement statistics are reported in Extended Data Table 2, while modeled proteins and rRNA are shown in Extended Data Table 3. Validation of the final models was done via MolProbity ^43^. Figures were generated using ChimeraX ^44^.

## Acknowledgments

All cryo-EM data used in this work were collected at the Karolinska Institutet’s 3D-EM facility. The SciLifeLab cryo-EM facility (used for grid preparation and initial screening) is funded by the Knut and Alice Wallenberg, Family Erling Persson, and Kempe foundations. We acknowledge the support of: Knut and Alice Wallenberg Foundation (KAW 2018.0080) to B.M.H and J.R.; the Swedish Research Council (2018-3808) to B.M.H.; Karolinska Institutet and the Max Planck Institute to J.R. J.R. is a Fellow of the Knut and Alice Wallenberg Foundation (WAF 2017).

## Author contributions

M.C. with A.K and J.R. help performed cell biology, biochemistry and sample preparations; G.V.G. and B.M.H. performed cryo-EM data collection; G.V.G. and B.M.H. processed data and determined structures; G.V.G. built and refined models with help from A.K. and B.M.H.; M.C. and G.V.G. wrote the manuscript with input from all authors; J.R. and B.M.H. oversaw the project and edited the manuscript.

## Competing interests

Authors declare no competing interests.

## Notes

### Competing Interest Statement

The authors have declared no competing interest.

