## Supplemental data for "Structural basis for late maturation steps of the human mitoribosomal large subunit"

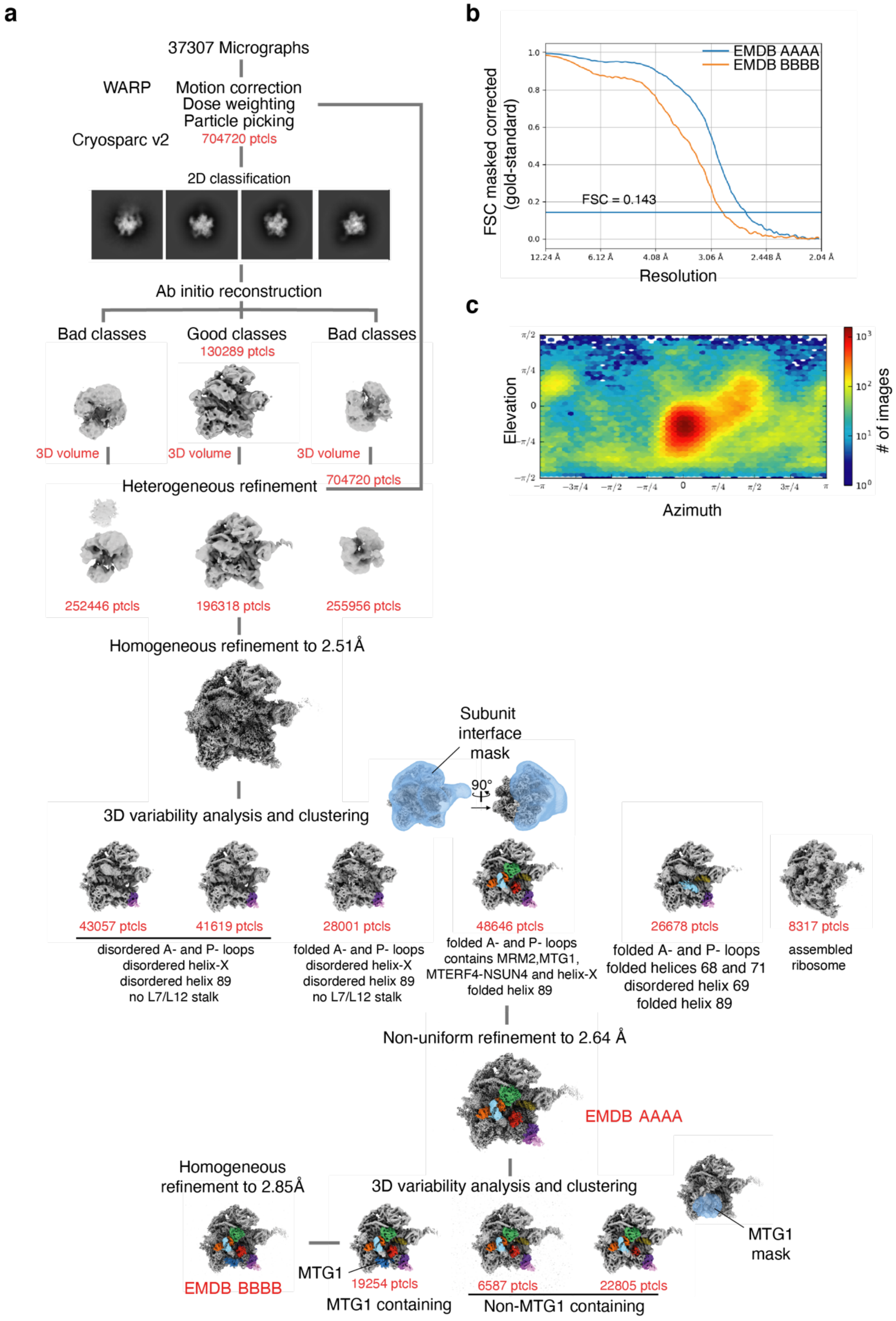


**Extended Data Fig. 1 Data processing strategy for the GTPBP5$^{KO}$ dataset.**
**a** Data processing strategy for the two deposited reconstructions (EMDB AAAA and BBBB) from the GTPBP5$^{KO}$ dataset. Colouring as in Figure 1. For the 3D variability analysis, the mask used is shown as a blue semi-transparent surface. **b** Gold-standard Fourier shell correlation (FSC) [45] for the two EMDB-deposited reconstructions. The horizontal blue line indicates the FSC cut-off at 0.143. **c**

Heatmap of the angular distribution for particle projections after heterogeneous refinement (196318 particles).

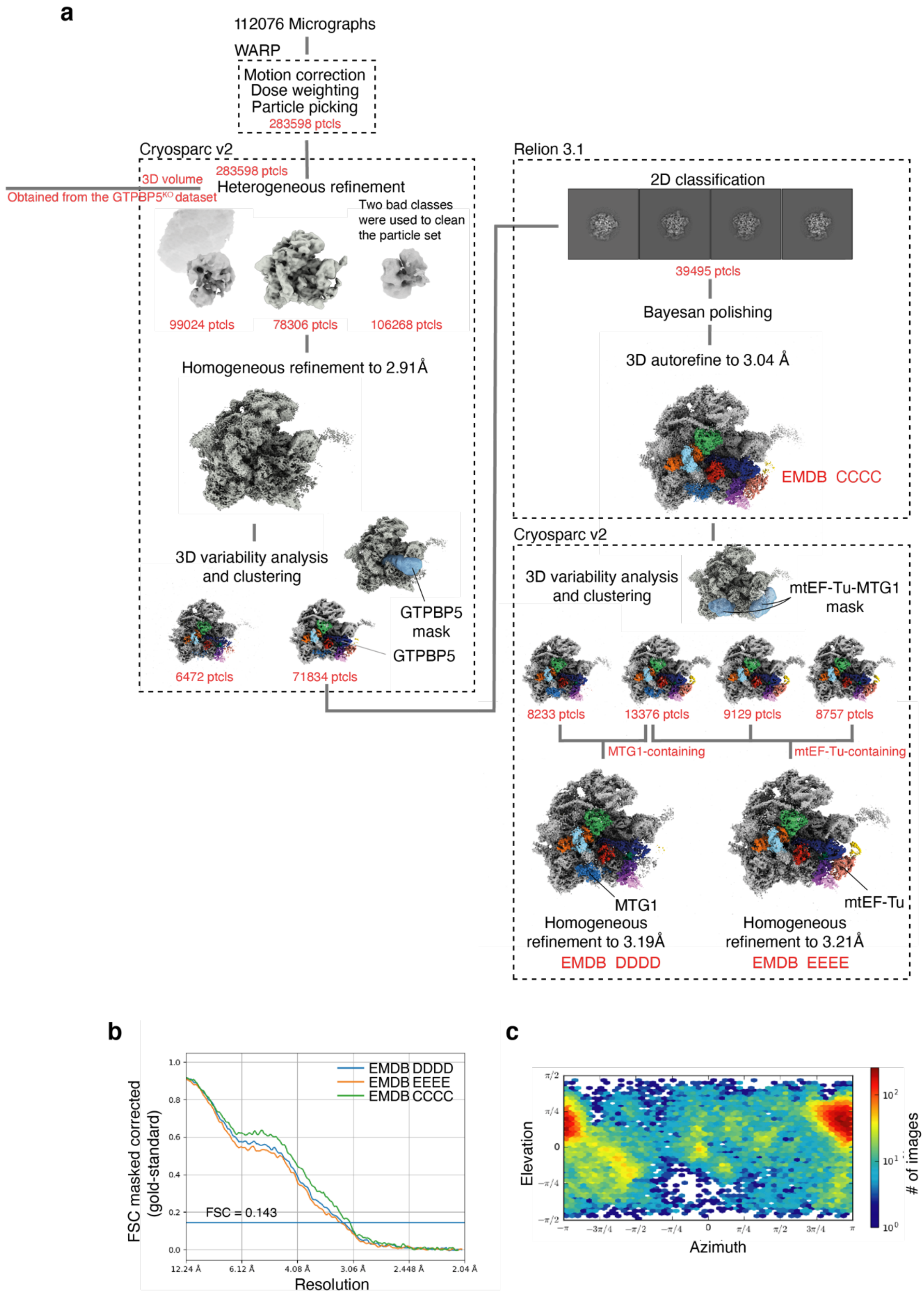


**Extended Data Fig. 2 Data processing strategy for the GTPBP5$^{IP}$ dataset.**

**a** Data processing strategy for the three deposited reconstructions (EMDB CCCC, DDDD and EEEE) from the GTPBP5$^{IP}$ dataset. Colouring as in Figure 1. For the 3D variability analysis, the mask used is shown as a blue semi-transparent surface. **b** Gold-standard Fourier shell correlation (FSC) [45] for the three EMDB-deposited reconstructions. The horizontal blue line indicates the FSC cut-off at

0.143. **c** Heatmap of the angular distribution for particle projections after homogeneous refinement (78306 particles).

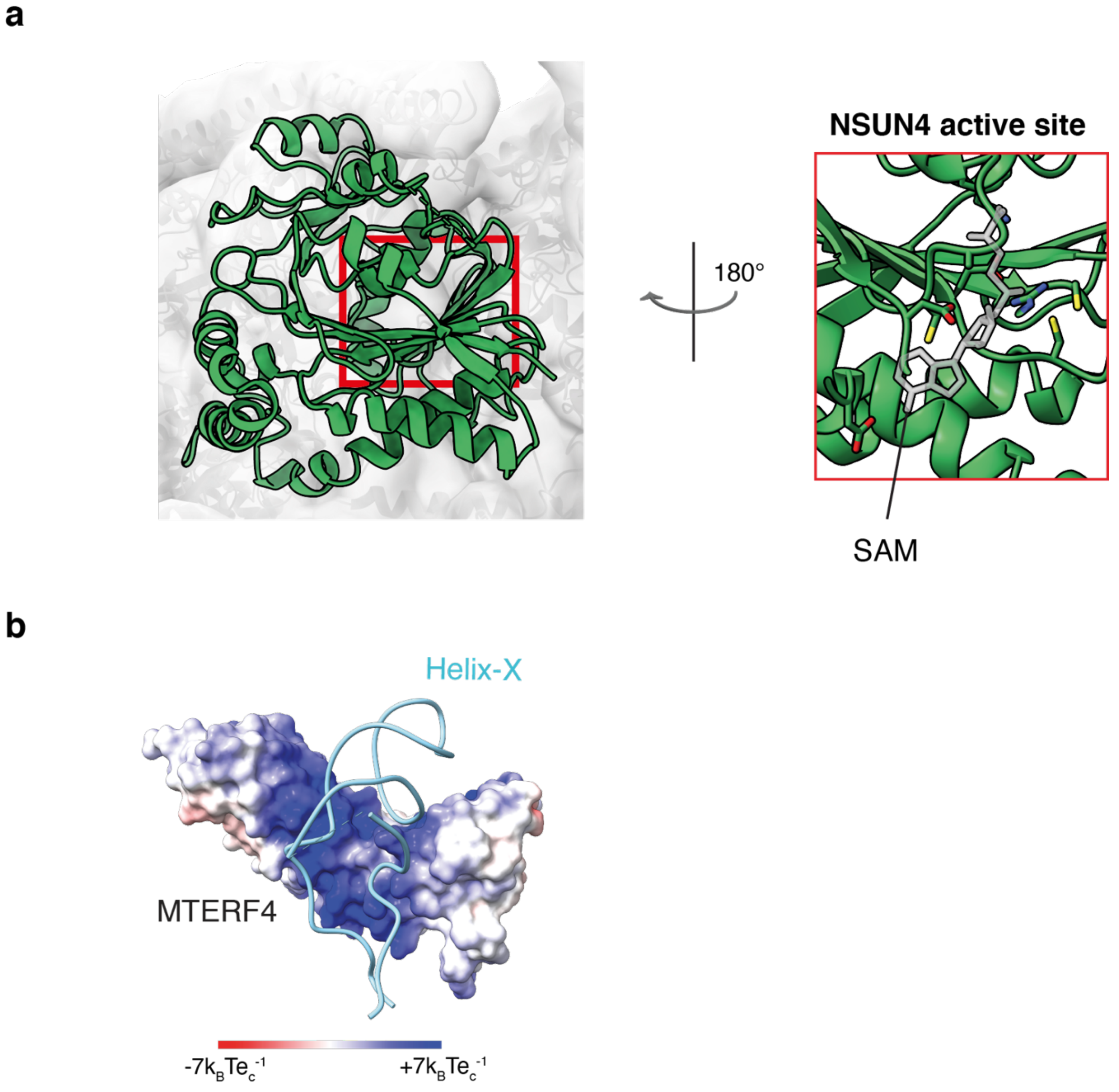


**Extended Data Fig. 3 NSUN4 active site and surface electrostatic potential of MTERF4.**
**a** View of NSUN4 in both GTPBP5$^{KO}$ and GTPBP5$^{IP}$ structures facing the mt-LSU. NSUN4 active site is indicated by a red rectangle. The right zoomed-in panel obtained after 180°C rotation in respect to the left panel shows the residues involved in SAM coordination and SAM as sticks. **b** Red and blue refer to electronegative and electropositive regions, respectively. ABPS was used to calculate the electrostatic potential [46].

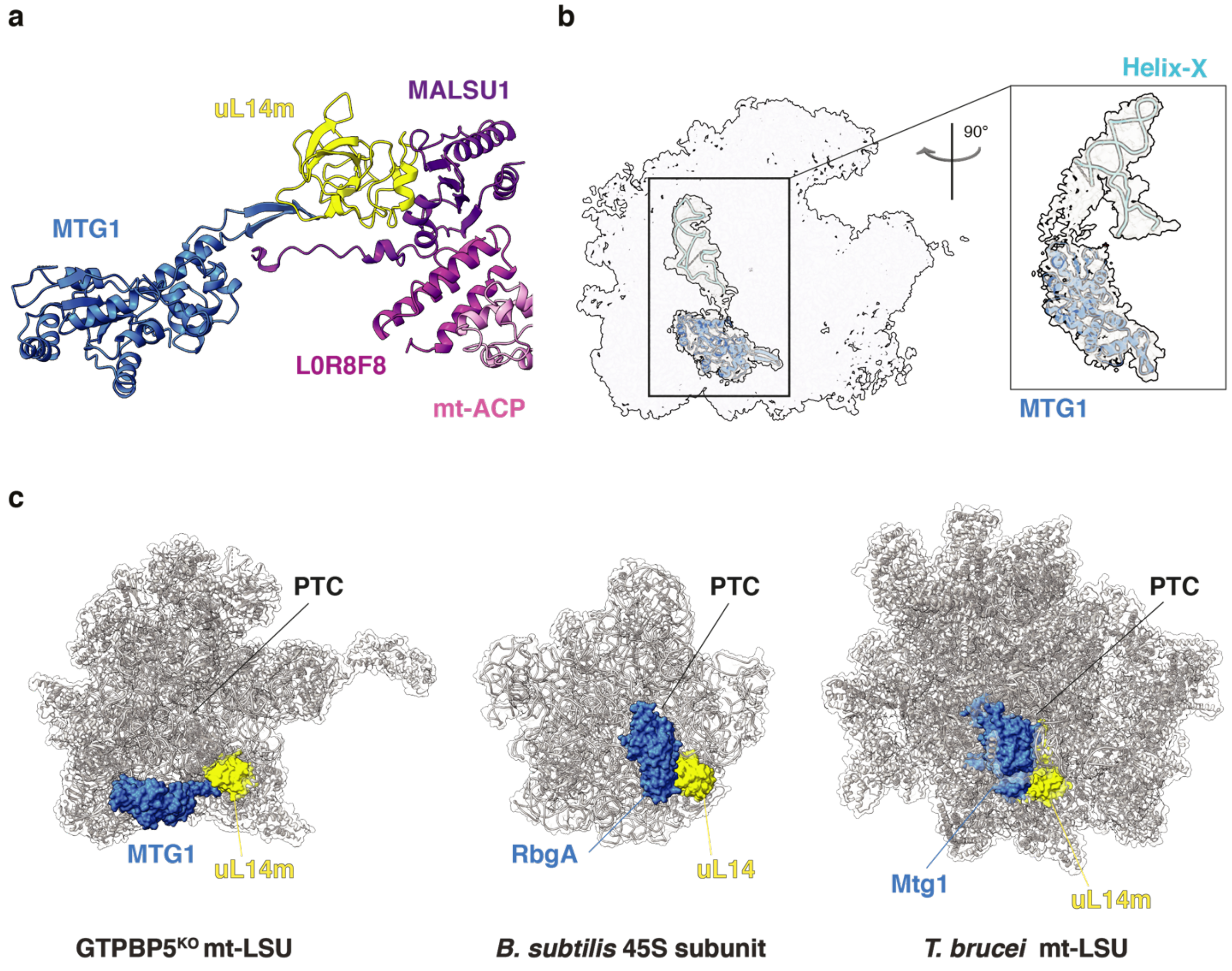


**Extended Data Fig. 4 Structural features of MTG1 and the MALSU1 module.**
**a** Cartoon representation of MALSU1 contacting MTG1. L0R8F8, mt-ACP and uL14m are also shown. **b** Cryo-EM density of the MTG1-helix-X contact on the $GTPBP5^{KO/IP}$ mt-LSU. The rotated zoomed-in panel shows the density of MTG1 contacted by the bottom part of Helix-X that was not modelled due to the lower resolution. Helix-X is represented in light blue while MTG1 is represented in pale blue. **c** Comparison of MTG1 position on the $GTPBP5^{KO/IP}$ mt-LSU, with its homologues RbgA on the *B. subtilis* 45S subunit (PDB: 6PPK [22]) and Mtg1 on the *T. brucei* mt-LSU (PDB: 6YXY [3]). Mitoribosomal protein uL14m that is situated in close proximity to MTG1 is coloured in yellow, while the PTC region is indicated with a label. MTG1, RbgA and Mtg1 are indicated in pale blue colour.

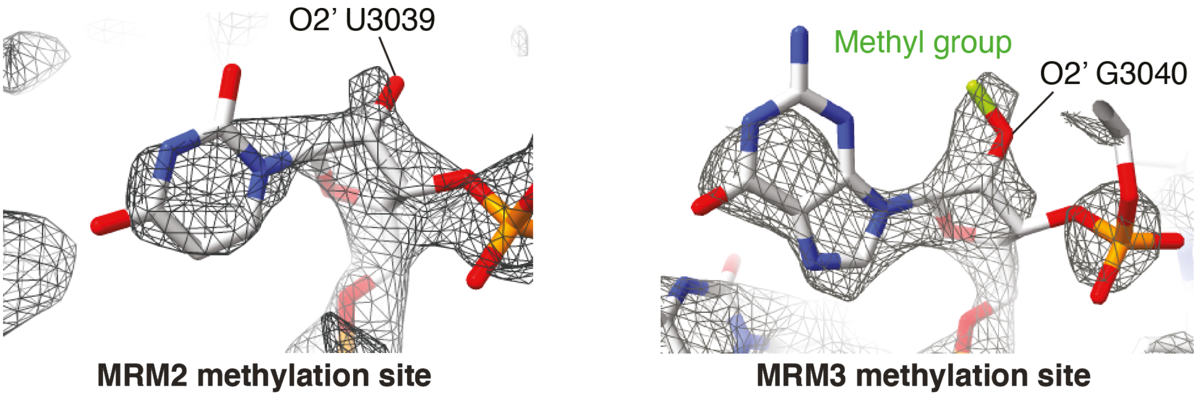


**Extended Data Fig. 5 MRM2 and MRM3 methylation sites on the mt-LSU.**
Representation of MRM2 (U3039; left) and MRM3 (G3040; right) methylation sites. The 2′-O of the ribose is highlighted in both views. The methyl group of MRM3 methylation site is indicated in green (right). The methyl group is missing in MRM2 methylation site (left).

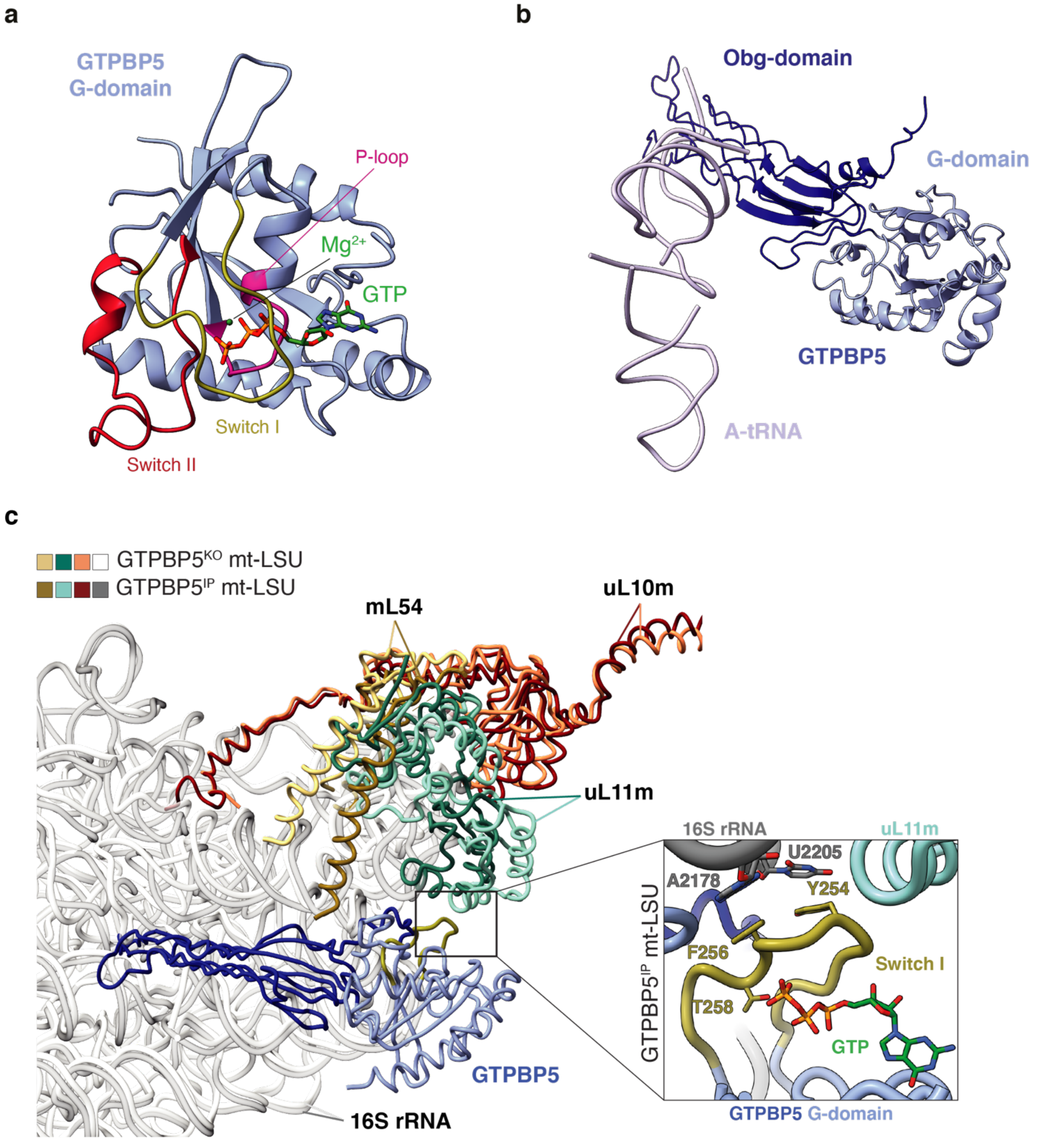


**Extended Data Fig. 6 Structural features of GTPBP5 and its interaction with the mt-LSU.**
**a** GTPBP5 G-domain displays GTP (green) in its binding pocket. Switch I, Switch II, the P-loop and the $Mg^{2+}$ are indicated. **b** Comparison of the $GTPBP5^{IP}$ structure with the translating mitoribosome (PDB: 5AJ4 [47]) shows that GTPBP5 overlaps with the A-site tRNA. **c** Representation of the L7/L12 stalk movement in $GTPBP5^{IP}$ mt-LSU when compared to the $GTPBP5^{KO}$ mt-LSU. The 16S mt-rRNA, mL54, uL10m and uL11m are highlighted in both structures. GTPBP5 is also shown. The right zoomed-in panel features interactions between GTPBP5 G-domain and the 16S mt-rRNA in $GTPBP5^{IP}$ mt-LSU structure. The G-domain Switch I and GTP are shown in yellow and green, respectively.

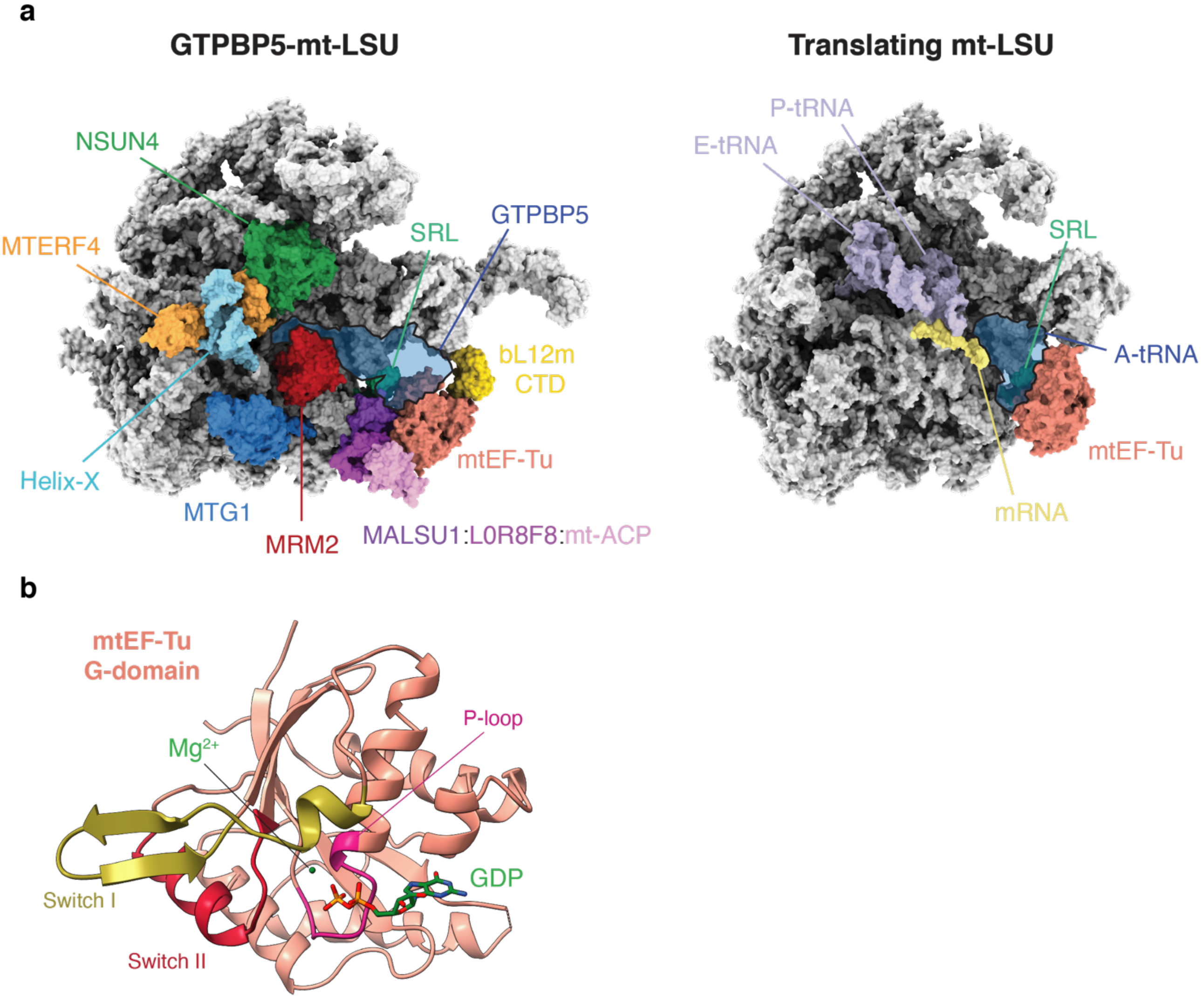


**Extended Data Fig. 7 mtEF-Tu interaction with the mt-LSU during mitoribosome assembly or translation and structural features of the G-domain.**

**a** Surface representation of the GTPBP5$^{IP}$ mt-LSU and the translating mt-LSU (PDB: 7A5G [27]) showing the different position of mtEF-TU. In GTPBP5$^{IP}$ mt-LSU structure, mtEF-Tu, MTERF4-NSUN4, MRM2, MTG1, GTPBP5, MALSU1 module, bL12 CTD, helix-X and SRL are indicated. In the translating mt-LSU structure the A-, P- and E-tRNAs are shown together with mtEF-Tu, SRL and mRNA (PDB: 7A5G [27]). GTPBP5 and the A-site tRNA in the respective structures are shown semi-trasparent. **b** mtEF-Tu G-domain displaying GDP (green) in its binding pocket. Switch I, Switch II, the P-loop and the $Mg^{2+}$ are indicated.

#### Extended Data Table 1 Cryo-EM data collection, refinement and validation statistics

| | GTPBP5$^{IP}$<br><br>(EMDB-CCCC)<br>(EMDB-DDDD for class with 100% MTG1 occupancy)<br>(EMDB-EEEE for class with 100% mtEF-TU occupancy)<br>(PDB-xxxx) | GTPBP5$^{KO}$<br><br>(EMDB-AAAA)<br>(EMDB-BBBB for class with 100% MTG1 occupancy)<br>(PDB-xxxx) |
|---|---|---|
| **Data collection and processing** | | |
| Microscope | FEI Titan Krios G3i | |
| Detector | K3 | |
| Voltage (kV) | 300 | |
| Electron exposure ($e^{-}$/Å$^2$/sec) | 49.265 | |
| Energy filter slit width (eV) | 10 | |
| Pixel size (Å) | 0.500 | |
| Magnification | 165000 | |
| Defocus range (µm) | -0.3 to -1.1 | |
| Map Resolution at FSC = 0.143 (Å) | 3.04 for EMDB-CCCC<br>3.19 for EMDB-DDDD<br>3.21 for EMDB-EEEE | 2.64 for EMDB-AAAA<br>2.85 for EMDB-BBBB |
| Sharpening B factor (Å$^2$) | -35.47 for EMDB-CCCC<br>-12.2 for EMDB-DDDD<br>-10.1 for EMDB-EEEE | -36.7 for EMDB-AAAA<br>-25.5 for EMDB-BBBB |
| **Refinement** | | |
| Initial model used (PDB code) | 5OOL | 5OOL |
| Model resolution at FSC = 0.143 (Å) | 3.08 | 2.63 |
| Model Composition | | |
| No. of chains | 74 | 68 |
| Total atoms | 115947 | 110267 |
| Protein residues | 10335 | 9606 |
| RNA residues | 1498 | 1515 |
| Ligands: GTP/GDP/SAM/SAH/PNS/$Mg^{2+}$/$Zn^{2+}$ | 1/2/1/1/1/93/2 | 0/1/1/0/1/112/2 |
| B Factors (min/max/mean) (Å$^2$) | | |
| Protein | 95.70/457.80/174.22 | 47.76/540.67/109.83 |
| RNA | 98.46/450.25/144.63 | 44.15/512.83/80.77 |
| Ligands | 86.04/317.83/213.67 | 40.16/191.65/130.73 |
| Validation | | |
| RMSD bonds (ouliers) (Å) | 0.004 (0) | 0.005 (6) |
| RMSD angles (outliers) (°) | 0.615 (11) | 0.631 (26) |
| Clashscore | 12.87 | 9.42 |
| MolProbity score | 1.97 | 2.02 |
| Rotamer outliers (%) | 0.000 | 2.53 |
| Ramachandran plot | | |
| Favored/Allowed/Outliers (%) | 94.99/4.95/0.06 | 96.60/3.35/0.05 |
| RNA Validation | | |
| Sugar pucker outliers (%) | 1.67 | 1.65 |
| Angle/bond outliers (%) | 0.1 | 0 |
| Bond outliers (%) | 0 | 0 |

EMDB, Electron Microscopy Data Bank; PDB, Protein Data Bank; RMSD, root-mean-square deviation; FSC, Fourier shell correlation.
GTP, guanosine triphosphate; GDP, guanosine diphosphate; SAH, S-adenosylhomocysteine; SAM, S-adenosyl methionine;
PNS 4'-phosphopantetheine

### Extended Data Table 2 Protein and RNA components of the mt-LSU assembly intermediates.

| Chain | Modelled residues GTPBP5$^{KO}$ | Modelled residues GTPBP5$^{IP}$ | UniprotID | Description | short name |
|---|---|---|---|---|---|
| 0 | 79-200 | 79-200 | Q9BYC8 | 39S ribosomal protein L32, mitochondrial, L32mt, MRP-L32 | RM32_HUMAN |
| 1 | 14-65 | 14-65 | O75394 | 39S ribosomal protein L33, mitochondrial, L33mt, MRP-L33 | RM33_HUMAN |
| 2 | 48-92 | 48-92 | Q9BQ48 | 39S ribosomal protein L34, mitochondrial, L34mt, MRP-L34 | RM34_HUMAN |
| 3 | 94-188 | 94-188 | Q9NZE8 | 39S ribosomal protein L35, mitochondrial, L35mt, MRP-L35 | RM35_HUMAN |
| 4 | 66-200 | 66-103 | Q9P0J6 | 39S ribosomal protein L36, mitochondrial, L36mt, MRP-L36 | RM36_HUMAN |
| 5 | 31-422 | 31-422 | Q9BZE1 | 39S ribosomal protein L37, mitochondrial, L37mt, MRP-L37 | RM37_HUMAN |
| 6 | 27-79,99-209,213-282,291-380 | 27-79,99-209,213-380 | Q96DV4 | 39S ribosomal protein L38, mitochondrial, L38mt, MRP-L38 | RM38_HUMAN |
| 7 | 36-322 | 36-325 | Q9NYK5 | 39S ribosomal protein L39, mitochondrial, L39mt, MRP-L39 | RM39_HUMAN |
| 8 | 97-181 | 97-181 | Q9NQ50 | 39S ribosomal protein L40, mitochondrial, L40mt, MRP-L40 | RM40_HUMAN |
| 9 | 15-137 | 15-137 | Q8IXM3 | 39S ribosomal protein L41, mitochondrial, L41mt, MRP-L41 | RM41_HUMAN |
| A | 1671-3388 | 1671-3386 | NA | 16S RRNA | 16S RRNA |
| A1 | 36-108,123-384, SAM | 27-108,123-384, SAM | Q96CB9 | 5-methylcytosine rRNA methyltransferase NSUN4 | NSUN4_HUMAN |
| B | 1603-1670 | 1603-1670 | NA | MT-TRNAVAL | MT-TRNAVAL |
| A2 | 90-327 | 90-327 | Q7Z6M4 | Transcription termination factor 4, mitochondrial | MTEF4_HUMAN |
| C | 55-185,213-401, GDP | 55-185,213-401, GDP | Q9BT17 | Mitochondrial ribosome-associated GTPase 1 | MTG1_HUMAN |
| D | 61-401 | 61-401 | Q5T653 | 39S ribosomal protein L2, mitochondrial, L2mt, MRP-L2 | RM02_HUMAN |
| t1 | 46-92 | 46-92 | P52815 | 39S ribosomal protein L12, mitochondrial | RM12_HUMAN |
| E | 45-348 | 45-348 | P09001 | 39S ribosomal protein L3, mitochondrial, L3mt, MRP-L3 | RM03_HUMAN |
| t2 | 62-91 | 62-91 | P52815 | 39S ribosomal protein L12, mitochondrial | RM12_HUMAN |
| F | 45-294 | 45-294 | Q9BYD3 | 39S ribosomal protein L4, mitochondrial, L4mt, MRP-L4 | RM04_HUMAN |
| G | Not present | 59-402, GTP | Q9H4K7 | Mitochondrial ribosome-associated GTPase 2 (GTP-binding protein 5) | MTG2_HUMAN |
| H | 53-147 | 53-147 | Q9BYD2 | 39S ribosomal protein L9, mitochondrial, L9mt, MRP-L9 | RM09_HUMAN |
| I | 36-240 | 29-240 | Q7Z7H8 | 39S ribosomal protein L10, mitochondrial, L10mt, MRP-L10 | RM10_HUMAN |
| J | 18-157 | 18-157 | Q9Y3B7 | 39S ribosomal protein L11, mitochondrial, L11mt, MRP-L11 | RM11_HUMAN |
| K | 2-178 | 2-178 | Q9BYD1 | 39S ribosomal protein L13, mitochondrial, L13mt, MRP-L13 | RM13_HUMAN |
| L | 31-145 | 31-145 | Q6P1L8 | 39S ribosomal protein L14, mitochondrial, L14mt, MRP-L14 | RM14_HUMAN |
| M | 10-301 | 10-301 | Q9P015 | 39S ribosomal protein L15, mitochondrial, L15mt, MRP-L15 | RM15_HUMAN |
| N | 32-133,148-251 | 62-134,146-251 | Q9NX20 | 39S ribosomal protein L16, mitochondrial, L16mt, MRP-L16 | RM16_HUMAN |
| O | 9-160 | 9-160 | Q9NRX2 | 39S ribosomal protein L17, mitochondrial, L17mt, MRP-L17 | RM17_HUMAN |
| P | 39-179 | 39-179 | A8K9D2 | Mitochondrial ribosomal protein L18, isoform CRA_b | A8K9D2_HUMAN |
| Q | 74-290 | 74-290 | P49406 | 39S ribosomal protein L19, mitochondrial, L19mt, MRP-L19 | RM19_HUMAN |
| R | 10-149 | 10-149 | Q9BYC9 | 39S ribosomal protein L20, mitochondrial, L20mt, MRP-L20 | RM20_HUMAN |
| S | 49-204 | 49-204 | Q7Z2W9 | 39S ribosomal protein L21, mitochondrial, L21mt, MRP-L21 | RM21_HUMAN |
| T | 47-301 | 47-301 | E7ESL0 | 39S ribosomal protein L22, mitochondrial | E7ESL0_HUMAN |
| U | 2-112,126-153 | 2-112,126-153 | Q16540 | 39S ribosomal protein L23, mitochondrial, L23mt, MRP-L23 | RM23_HUMAN |
| V | 15-216 | 15-216 | Q96A35 | 39S ribosomal protein L24, mitochondrial, L24mt, MRP-L24 | RM24_HUMAN |
| W | 48-201 | 48-201 | Q9P0M9 | 39S ribosomal protein L27, mitochondrial, L27mt, MRP-L27 | RM27_HUMAN |
| X | 2-244 | 2-244 | Q13084 | 39S ribosomal protein L28, mitochondrial | RM28_HUMAN |
| Y | 63-238 | 63-238 | Q9HD33 | 39S ribosomal protein L47, mitochondrial, L47mt, MRP-L47 | RM47_HUMAN |
| Z | 35-154 | 35-154 | Q8TCC3 | 39S ribosomal protein L30, mitochondrial, L30mt, MRP-L30 | RM30_HUMAN |
| a | 35-77,104-142 | 35-77,104-142 | Q9Y6G3 | 39S ribosomal protein L42, mitochondrial, L42mt, MRP-L42 | RM42_HUMAN |
| b | 2-149 | 2-149 | Q8N983 | 39S ribosomal protein L43, mitochondrial, L43mt, MRP-L43 | RM43_HUMAN |
| c | 31-316 | 31-107,119-316 | Q9H9J2 | 39S ribosomal protein L44, mitochondrial, L44mt, MRP-L44 | RM44_HUMAN |
| d | 36-54,69-91,117-294 | 36-52,70-91,111-294 | Q9BRJ2 | 39S ribosomal protein L45, mitochondrial, L45mt, MRP-L45 | RM45_HUMAN |
| f | 48-66,77-132,150-212 | 48-66,77-132,150-193,196-212 | Q96GC5 | 39S ribosomal protein L48, mitochondrial, L48mt, MRP-L48 | RM48_HUMAN |
| e | 43-104,116-217,227-279 | 43-104,116-217,227-279 | Q9H2W6 | 39S ribosomal protein L46, mitochondrial, L46mt, MRP-L46 | RM46_HUMAN |
| g | 36-201 | 38-201 | Q13405 | 39S ribosomal protein L49, mitochondrial, L49mt, MRP-L49 | RM49_HUMAN |
| h | 51-78,82-158 | 52-158 | Q8N5N7 | 39S ribosomal protein L50, mitochondrial, L50mt, MRP-L50 | RM50_HUMAN |
| i | 32-128 | 32-128 | Q4U2R6 | 39S ribosomal protein L51, mitochondrial, L51mt, MRP-L51 | RM51_HUMAN |
| j | 24-108 | 24-108 | A8K7J6 | 39S ribosomal protein L52, mitochondrial | A8K7J6_HUMAN |
| k | 2-96 | 13-56,61-96 | Q96EL3 | 39S ribosomal protein L53, mitochondrial, L53mt, MRP-L53 | RM53_HUMAN |
| l | 1-26,94-136 | 1-26,94-136 | Q6P161 | 39S ribosomal protein L54, mitochondrial | RM54_HUMAN |
| m | 34-78 | 34-78 | Q7Z7F7 | 39S ribosomal protein L55, mitochondrial | RM55_HUMAN |
| n | 25-239 | 51-239, SAH | Q9UI43 | rRNA methyltransferase 2, mitochondrial (Homo sapiens) | MRM2_HUMAN |
| p | 38-61,70-83,95-163,174-193 | 38-61,70-83,95-163,174-193 | Q14197 | Peptidyl-tRNA hydrolase ICT1, mitochondrial, EC 3.1.1.29 | ICT1_HUMAN |
| o | 23-102 | 12-102 | Q9BQC6 | Ribosomal protein 63, mitochondrial, hMRP63 | RT63_HUMAN |
| q | 25-159 | 25-159 | Q8TAE8 | Growth arrest and DNA damage-inducible proteins-interacting protein 1 | G45IP_HUMAN |
| r | 35-41,47-196 | 35-41,47-196 | Q9NVS2 | 39S ribosomal protein S18a, mitochondrial, MRP-S18-a, Mrps18a | RT18A_HUMAN |
| s | 41-119,138-430 | 41-119,140-430 | Q9NP92 | 39S ribosomal protein S30, mitochondrial, MRP-S30, S30mt | RT30_HUMAN |
| t | Not present | 55-184,190-440, GDP | P49411 | Elongation factor Tu, mitochondrial | EFTU_HUMAN |
| u | 91-219 | 87-219 | Q96EH3 | Mitochondrial assembly of ribosomal large subunit protein 1 (Homo sapiens) | MASU1_HUMAN |
| v | 2-70, PNS | 2-70, PNS | L0R8F8 | MIEF1 upstream open reading frame protein (Homo sapiens) | MIDUO_HUMAN |
| w | 70-156 | 70-156 | O14561 | Acyl carrier protein, mitochondrial (Homo sapiens) | ACPM_HUMAN |
| t3 | 62-91 | 62-91 | P52815 | 39S ribosomal protein L12, mitochondrial | RM12_HUMAN |
| t4 | 62-90 | 62-90 | P52815 | 39S ribosomal protein L12, mitochondrial | RM12_HUMAN |
| t5 | 62-91 | 62-91 | P52815 | 39S ribosomal protein L12, mitochondrial | RM12_HUMAN |
| t6 | 63-89 | 63-89 | P52815 | 39S ribosomal protein L12, mitochondrial | RM12_HUMAN |
| t7 | Not present | 129-198 | P52815 | 39S ribosomal protein L12, mitochondrial | RM12_HUMAN |

GTP, guanosine triphosphate; GDP, guanosine diphosphate; SAH, S -adenosylhomocysteine; SAM, S-adenosyl methionine; PNS 4'-phosphopantetheine
